# Reversible coating of cells with synthetic polymers for mechanochemical regulation of cell adhesion

**DOI:** 10.1101/2020.10.25.354480

**Authors:** Yoshihisa Kaizuka, Rika Machida

**Author notes:** Send correspondence to Yoshihisa Kaizuka.

## Abstract

The chemical control of cell–cell interactions using synthetic materials is useful for a wide range of biomedical applications. Herein, we report a method to regulate cell adhesion and dispersion by introducing repulsive forces to live cell membranes. To induce repulsion, we tethered amphiphilic polymers, such as cholesterol-modified polyethylene glycol (PEG-CLS) to cell membranes. These amphiphilic polymers both bind to and dissociate rapidly from membranes and thus, enable the reversible coating of cells by mixing and washout without requiring genetic manipulation or chemical synthesis in the cells. We found that the repulsive forces introduced by these tethered polymers can induce cell detachment from a substrate and allow cell dispersion in a suspension, modulate the speed of cell migration, and improve the separation of cells from tissues. Our analyses showed that coating the cells with tethered polymers most likely generated two distinct repulsive forces, lateral tension and steric repulsion, on the surface, which can be tuned by altering the polymer size and density. We also modeled how these two forces can be generated in kinetically distinctive manners to explain the various responses of cells to the coating. Collectively, our observations and analyses show how we can mechanochemically regulate cell adhesion and dispersion and may contribute to the optimization of chemical coating strategies for regulating various types of cell–cell interacting systems.

## Introduction

Cell adhesion, interaction, and dispersion regulate a wide range of *in vitro* and in vivo processes ^1–6^. Both 2D and 3D *in vitro* cell culture requires the control of cell–cell or cell–substrate interactions. The processes involved in collecting cells from tissues or preparing cells for flow cytometry analysis and single cell genetics require cell dispersion at a single cell level. The adsorption of cells to the surface of biomaterials needs to be managed, and cell–cell interactions and cellular communications are essential for the formation of spheroids and organoids. Processes *in vivo* involve more complicated cell–cell interactions, and the fine-tuning of such cell–cell communication has enabled cancer immunology therapies using checkpoint inhibitors. The cell–cell interface involves many binding proteins and the extracellular matrix, and the mechanical properties of the cell membranes and the cytoskeleton can also modulate cell–cell interactions ^7^.

The precise control of cell adhesion and dispersion with synthetic materials is a promising approach for future therapeutics, but is still a challenge to achieve ^8–9^. For example, multiple strategies involving modifications of the plasma membrane of cells have been developed but each method has its advantages and disadvantages. DNA hybridization has been used for controlling specific cell–cell interactions, but the applications are limited because of difficulties in molecular control and stability ^10^. We have found that polymers modified with multiple cholesterol moieties can bind to cell membranes and function as adhesive molecules ^11^. However, the cell adhesion mediated by such an amphiphilic polymer was very different from the cell adhesion mediated by naturally expressed adhesive proteins. In addition, these polymers restricted the spreading of the area of cell adhesion; thus, effectively induced a repulsive force on the cell membrane, indicating the intrinsic difficulty of artificially achieving cell adhesion by modifying cell membranes. However, this chemically induced repulsive force exerted by the polymers on the cell membranes suggested that this system can potentially be used for cell dispersion, which could be an alternative to the current cell dispersion methods that involve partial and irreversible disruption of samples either mechanically or enzymatically.

Herein, we report a method for the chemical control of cell adhesion and dispersion. By introducing repulsive forces on cell membranes using amphiphilic polymers, we succeeded in the chemical control of cells. This method does not involve any genetic manipulation, fabrication of substrates, or chemical synthesis involving the cells, but the synthetic cholesterol-modified amphiphilic polymers were simply added to the cells. These amphiphilic polymers, which had good water solubility and a high binding rate constant (k_on_) with lipid membranes also had a high dissociation rate constant (k_off_); thus, these polymers can be easily removed from lipid membranes by washout. These binding and dissociation cycles can enable the reversible and temporary coating of cell membranes, resulting in reduced damage to cells. We show that coating cells with these amphiphilic polymers at a density of only a fraction of the total density of the surface-membrane expressed proteins could modulate the adhesion and dispersion of cells.

## Experimental sections

### Cells and chemicals

HEK 293, HeLa, Jurkat, and NIH-3T3 cells were obtained from JCRB Cell Bank (Osaka, Japan). MC 38 and K562 cells were obtained from the Cell Resource Center for Biomedical Research, Institute of Development, Aging and Cancer, Tohoku University. PEG-CLS was purchased from Biochempeg (Watertown, MA, USA). BSA-CLS was created by bioconjugation of BSA (Merck, Tokyo, Japan) and NHS-PEG1K-cholesterol (Nanocs, New York, NY, USA). Gelatin-CLS was synthesized as previously described ^11^. Calcein-AM and Cell Counting Kit-8 were purchased from Dojindo (Kumamoto, Japan).

### *In vitro* assays

Cell proliferation was measured using a CCK-8 kit. Cell morphology was measured by image analysis using ImageJ (NIH). In the cell-detachment assay, cells were placed in media that contained the indicated concentrations of polymers and were imaged. This assay was dependent on the cell culture and adhesion states. Thus, for experiments to be directly compared, we used cells cultured in a 4-well glass-bottom dish in which all the wells shared the same substrate (cover glass coated with adhesive polymers) (CELLview, Greiner, Tokyo, Japan). The area of cell adhesion that appeared dark in the RICM images was measured by ImageJ. In the cell-dispersion assay, intrinsically aggregating K562 cells in media were thoroughly pipetted once to disperse the cells, and then were mixed with the coating polymers and the cells were left for 1 h before images were taken. Media containing PEG-CLS was replaced with fresh media, and an image was taken again 2 h after the washout. The number of cells in clusters was counted from the images. Cell-migration assays were performed using a 2-well silicone insert with a defined cell-free gap (Ibidi, Gräfelfing, Germany). The polymers were added to the cell culture media immediately after the release of the silicone insert to induce cell migration, and the polymers were kept in the media during the experiment. Fluorescence, brightfield, and reflection interference microscope imaging were performed using a Leica (Solms, Germany) AF6000LX microscope equipped with a 100× 1.46 NA oil immersion objective and a Cascade II EMCCD camera (Roper, Tuscon, AZ, USA). For RICM imaging, a filter cube consisting of a narrowband-path filter (543–553 nm) and a beam splitter was used with arc lamp illumination.

### Animal experiments

Research involving animals complied with the protocols approved by the Animal Care and Use Committee of National Institute for Materials Science. Age-matched, 6–8 week old female C57BL/6 mice (Japan SLC. Hamamatsu, Japan) were implanted subcutaneously on the right flank with ~10^5^ of MC-38 tumor cells. After isolation, the grown tumors were dissected, divided, and then incubated in RPMI media that contained polymers or enzymes. After the tumors were incubated in the media at 37 °C for 30 min with occasional vortexing, the solution was centrifuged at 5 × *g* for 10 min, twice, to remove debris. The suspension was passed through a 70-μm cell-strainer. After further centrifugation, red blood cell lysis buffer (Bay Bioscience, Kobe, Japan) was added to cells for 3 min, and then washed out by further centrifugation. Cells were resuspended in RPMI buffer and were subjected to antibody staining with anti-CD45 (FITC, catalog no. 130-102-997), anti-CD11b (PE, catalog no. 130-113-806), and anti-CD11c (FITC, catalog no. 130-102-798) (Miltenyibiotec Tokyo, Japan). Stained cells were analyzed using Cell sorter SH800 (Sony, Tokyo, Japan).

## Results and Discussion

We evaluated the binding of cholesterol-modified polyethylene glycol (PEG-CLS), as a model amphiphilic coating material, to live cell membranes. PEG-CLS could be incorporated in the membranes of HEK 293 cells, as has been previously shown for cholesterol-modified gelatin ^11^, while unmodified soluble PEG methyl ether (mPEG) barely interacted with the cell surfaces (Figure 1A). Incorporation of PEG-CLS in the cell membranes was quantified by flow cytometry (Figure 1B). The molecular density of the tethered PEG-CLS in the cell membranes was calibrated as previously reported ^11^, and we found that the cell membranes contained 5000–10000 PEG-CLS molecules per μm^2^ when the cells were incubated with 10–100 μM PEG-CLS. The density of the bound PEG-CLS was independent of the size of PEG molecules, and was similar to the density of gel-CLS incorporated in cell membranes (2500–7500/μm^2^) ^11^. This density range was approximately 12%–33% of that of the membrane proteins natively expressed in live cell membranes (30000–40000/μm^2^) ^12^, and corresponded to < 0.7% of the total phospholipids packed in a 1-μm^2^ area. Thus, the exclusion volume created by the tethered PEG-CLS molecules can be altered, depending on the tethered density and size of the PEG molecules, but was still comparable to, or less than, the exclusion volume created by native membrane proteins that are expressed in a several-fold higher density.

**Figure 1.**
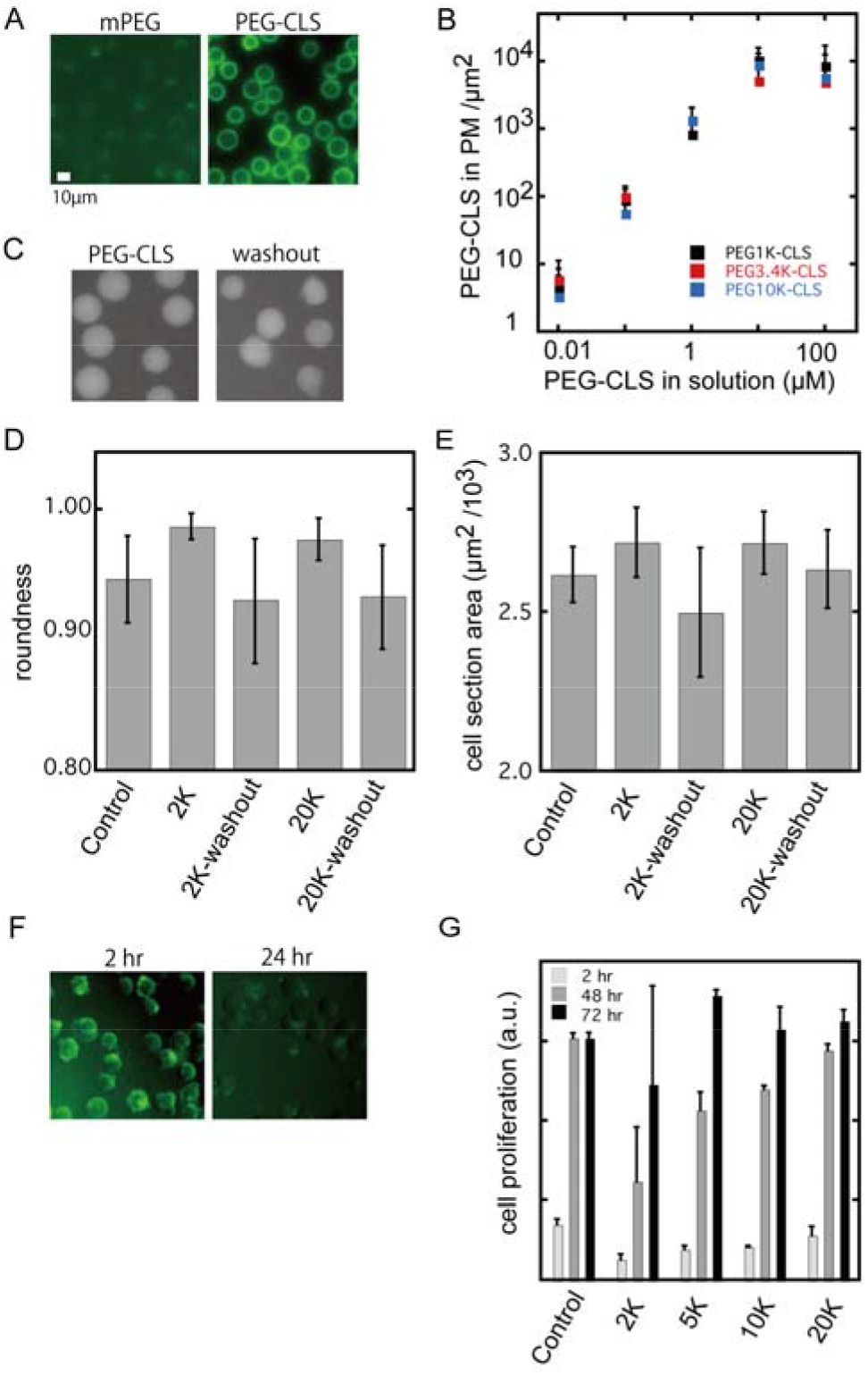
Coating of the cells with PEG-CLS. (A and B) Imaging and flow cytometry analysis of HEK 293 cells coated with PEG-CLS. HEK 293 cells (1.5 × 106/mL) in suspension were stained for 1 h with either 25 μM of 5K-mPEG and 5K-PEG-CLS containing 10% FITC-labeled molecules (A) or with a range of PEG-CLS molecules with various PEG sizes and various concentrations containing 5% FITC-labeled molecules (B), and were imaged or analyzed after washing once with PBS. (C–E) Calcein-AM-loaded HEK 293 cells uncoated (not shown) or coated with PEG-CLS (C, left), and at 2 h after washout of PEG-CLS (C, right) were observed, and the roundness (D) and volume (E) of the cells were calculated by image analyses. (F) Images of PEG-CLS-FITC in HEK 293 cells immediately after one washout (left) and 24 h later immediately after another washout. (G) Cell proliferation assay of HEK 293 cells treated with various sizes of PEG-CLS molecules for 2 h, immediately after the incubation and washout, and after culture for 48 and 72 h. Error bars represent standard deviations (B, G) or standard error of mean (D, E).

When coating the cell membranes, the amphiphilic polymers were tethered to lipid bilayers via cholesterol; thus, these polymers may facilitate the creation of a polymer brush structure on the membrane surface, potentially in combination with natively expressed proteins ^13–14^. Compared with highly densely packed polymers that are grafted on the surface and form an ideal true brush regime, such as was the case with > 0.065 chains/nm^2^ of 17000 Mw polyacrylamide ^15^, the density of the tethered PEG-CLS molecules on the cell membranes was only 7%–15%. More specifically, measurement of the PEG on a supported lipid bilayer showed that the transition from a mushroom regime to a brush regime occurred at a density of 4%–8% of PEG lipids in the membranes, which was significantly higher than the observed 0.7% incorporation of PEG-CLS in cells ^16^. The conventional model, Ng^6/5^>1/*ρ* _g_, also predicted that the measured density of PEG-CLS in our systems was 1–3 orders of magnitudes less than the density required for making a true brush of Mw 1000–20000 PEG molecules ^17^. Therefore, the tethered PEG-CLS molecules in the cell membranes are likely to form a mushroom regime and behave as two-dimensionally fluid molecules in the lipid bilayers without gelation ^16, 18–19^. Such fluidity is a feature of polymers tethered to lipid bilayers, which is very different from fixed polymers grafted on a substrate. The tethered polymers are inserted in the lipid bilayers and thus, may alter the molecular packing as well as the three-dimensional elasticity of the membranes.

The effect of the incorporation of amphiphilic polymers in the cell membranes was further analyzed. We observed the swelling of cells after coating with PEG-CLS (Figure 1C). There was a consistent increase in the roundness of cells coated with a saturated amount of either 2000 or 20000 molecular weight (Mw) PEG-CLS (Figure 1D). We also observed an increase in the volume of coated cells (Figure 1E). Upon swelling, the plasma membranes of coated cells were stretched and most likely tensioned more strongly, compared with those in uncoated cells. Such changes can be explained by the classical theory of membrane spontaneous curvature ^20–21^. Membranes have a natural curvature that results in a free energy minimum, and modulation of this spontaneous curvature has been observed with various examples of macromolecular insertion into lipid bilayers ^20, 22–24^. In particular, such artificial changes in curvature generate lateral tension ^25^. The drastic structural transition of budding has even been observed in highly tensioned lipid bilayers, and also in cells expressing a large amount of highly bulky glycocalyx^20, 22–24, 26^. The expansion of packed lipids in the bilayers has been observed with PEG lipid insertion using electron spin resonance and this may be the reason for the increase in the membrane spontaneous curvature ^27^. In the present study, the budding of cell membranes was not observed, probably because the cell membranes initially had a low volume to area ratio and could accommodate a large increase in membrane tension or because the cell membranes had sufficient stiffness, supported by the cytoskeleton and the bound molecules, to resist budding. Such increases in membrane bending rigidity and stiffness have been observed in lipid bilayers and cells bound with PEG or proteins ^28–29^. In addition, a membrane-condensing effect that is possibly induced by inserted cholesterol may also play a role ^30^, and the cells coated with PEG-CLS may actively adjust the cellular volumes in response to the spontaneous curvature change.

Importantly, we observed a rapid reversal of the cell swelling by decreasing the amount of PEG-CLS by washout in 2 h (Figure 1C-E). This observation indicated that reversible coating was possible, by taking advantage of the high k_off_ value of the amphiphilic polymers detaching from the membranes ^11^. Indeed, after culture for 24 h followed by another washout process, the level of retained PEG-CLS in the plasma membranes was significantly decreased (Figure 1F). We also investigated the toxicity of this reversible coating. It has been previously reported that culturing adherent cells with 2K-PEG-CLS at concentrations < 50 μM resulted in only a small effect on the cell viability ^31^. Here, we evaluated the toxicity of a suspension of cells, where the interaction of PEG-CLS and the cell membranes can be more rapid than that in an adherent culture, and only partial toxicity with less than an approximately10% drop in viability was observed over a short period of culture (1 h) at a relatively high concentration (25 μM) of PEG-CLS (data not shown). Suspension for longer than 2 h caused a severer effect (~30% loss) for shorter PEG molecules (2K), while the effect was still minimal for longer PEG molecules (20K). Such differential effects may reflect the stronger tendency of shorter PEG lipids to induce micelle formation ^27^. After washout of PEG-CLS, the cells that had been coated grew as rapidly as non-treated cells suggesting that the coating did not cause permanent damage to the cells (Figure 1G).

Previous observations have indicated that the insertion of cholesterol-modified polymers induced a repulsive interaction between cells and opposing surfaces ^11^. Here, we evaluated this repulsive force by measuring the adhesion of cells coated with PEG-CLS molecules and other amphiphilic polymers. Using time-lapse imaging of cell adhesion to the substrate by bright field and reflection interference contrast microscopy (BF and RICM), we observed the rapid induction of the detachment of HEK 293 cells coated with 50 μM of 3.4K-PEG-CLS, 5 μM gel-CLS, and 5 μM cholesterol-modified bovine serum albumin (BSA-CLS) (Figure 2A-B). Addition of unmodified polymers did not alter the adhesion state. The area of adhesion was measured from the RICM images where the adhered area appeared dark ^11^ (Figure 2A). Although incubated at a 10-fold lower concentration, the larger gel-CLS and BSA-CLS molecules induced a more rapid detachment than did PEG-CLS. This result suggested that larger sized molecules had a stronger effect.

**Figure 2.**
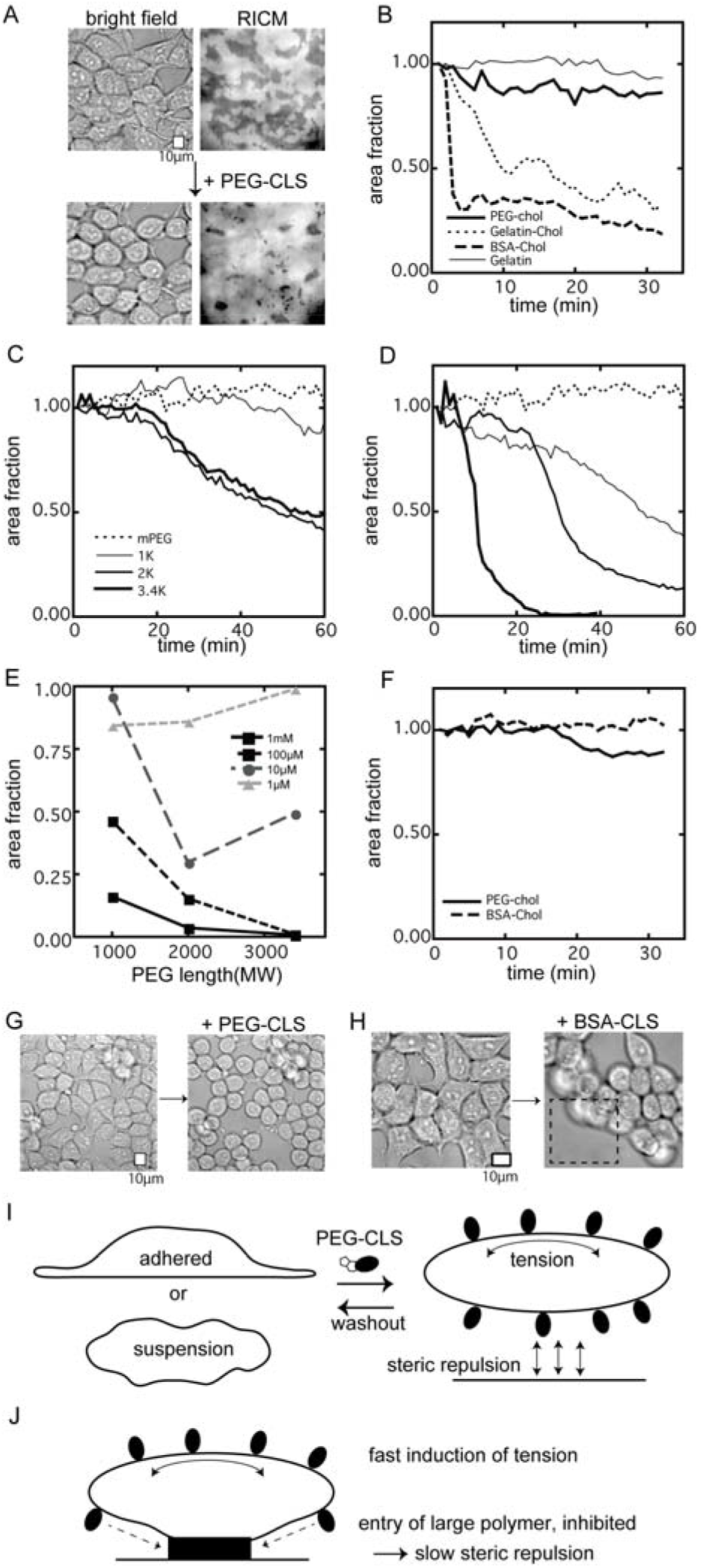
Detachment of coated cells from substrate. (A) Examples of the cell-detachment assay. HEK 293 cells adhered to the substrate were gradually detached after coating with PEG-CLS, and the area of adhesion that appeared dark in the RICM images was measured by image analysis. (B) Time course of the normalized adhesion area of HEK 293 cells treated with various types of amphiphilic polymers (50 μM 3.4K-PEG-CLS, 5 μM gelatin-cholesterol, 5 μM BSA-cholesterol, and 5 μM control gelatin without modification). (C and D) Time course of the normalized adhesion area of HEK 293 cells coated with 10 μM (C) or 100 μM (D) of different sized PEG-CLS molecules. (E) Normalized adhesion area of HEK 293 cells 60 min after coating with different concentrations of PEG-CLS molecules of various sizes. (F) Time course of the normalized adhesion area of HeLa cells treated with 50 μM 3.4K-PEG-CLS and 5 μM BSA-cholesterol. (G and H) Bright field images of HEK 293 cells treated with 50 μM 3.4K-PEG-CLS (G) and 5 μM BSA-cholesterol (H) at 0 and 30 min. (I) A model of how two distinct repulsive forces are generated in the cells coated with amphiphilic polymers. Cells in suspension and adherent cells could experience both membrane lateral tension and surface steric repulsion. (J) A model of the different kinetics between tension and steric repulsion.

The analysis of cells coated with PEG-CLS indicated that there were two distinct repulsive forces generated by the tethered polymers, namely steric repulsion and lateral tension (Figure 1). The mechanisms of cell detachment reflect how these polymers generated repulsive forces. Steric repulsion is caused by the polymer exclusion volume; thus, its strength depends on both the polymer size and the density of the tethered polymers However, cell swelling was observed that was independent of the size of the PEG molecules, as shown in Figure 1D, which suggested that the membrane tension was governed primarily by the density of the tethered polymers. To investigate in more detail how the cells respond to these forces, we compared the detachment of PEG-CLS-coated HEK 293 cells using various sizes of PEG molecules, 1–3.4K (average values). The PEG molecules in this Mw range correspond to 1.9–3.9 nm in Flory length ^32^, and thus, the exclusion volumes of 1K and 3.4K PEG molecules differ by approximately 8-fold ^29^. We observed that cells coated with PEG-CLS molecules of various sizes and at different concentrations detached with distinctive kinetics. For example, at a concentration of 10 μM, we observed a significant PEG size-dependent difference in the kinetics of cell detachment (Figure 2C–E). In contrast, at concentrations higher than 100 μM, cells coated with even short-chain PEG molecules detached rapidly (Figure 2D–E), and at a lower concentration of 1 μM, even long-chain PEG molecules were not effective in inducing detachment after 60 min (Figure 2E), suggesting PEG density-dependent regulation. Short-chain 1K PEG will still create an exclusion volume and steric repulsive force; thus, steric repulsion and tension were not completely decoupled in our experimental system. However, our observations suggested that the membrane tension primarily regulated the detachment of HEK 293 cells from a substrate over a wide range of PEG-CLS concentrations, while at the middle of the tested PEG-CLS concentration range, steric repulsion also becomes a critical factor in determining the kinetics of detachment.

We also tested the detachment of HeLa cells. Interestingly, the HeLa cells responded more strongly to a high concentration of PEG-CLS than to a low concentration of BSA-CLS, contrary to the results for HEK 293 cells (Figure 2A and 2F). These results suggested that, at least over a short time range of 30–60 min, the strong steric repulsion created by the large BSA molecules was not as effective in HeLa cells as in HEK 293 cells. In contrast, the membrane tension created by a high concentration of PEG-CLS resulted in the detachment of HeLa and HEK 293 cells at a similar rate.

Our observations suggested that both tension and steric repulsion contributed to the cell detachment but the effects can be slightly different in different adhesion structures. Tension is an effective force for the detachment of all types of adhesions, including specialized adhesive structures, such as focal adhesions, which are formed more in HeLa cells than in HEK 293 cells. In contrast, steric repulsion is also effective in all types of adhesions but depends more on the kinetics than does tension. Steric repulsion must function locally, thus, the large polymers need to be distributed to the site of adhesion. However, the access of such large molecules to the adhesion sites is likely strongly obstructed in tight and condensed junctions, such as focal adhesions. In contrast, membrane tension functions more globally, depending primarily on the total number of amphiphilic polymers tethered to a cell. These kinetic differences in how the two forces are generated may explain the different responses of the two cell lines. In another example, in HEK 293 cells, the gradual detachment of individual cells induced by PEG-CLS indicated that tension acted to detach cells from both the surfaces and neighboring cells (Figure 2G). In contrast, BSA-CLS frequently induced the abrupt detachment of a group of HEK 293 cells (Figure 2H), which may reflect the more rapid entry of the large BSA molecules and the rapid induction of steric repulsion at the cell–substrate interface rather than the cell–cell interface. The two types of repulsive forces acting on the cell membranes are summarized in Figure 2I and J.

We also evaluated the induction of the repulsive forces between cells for use in cell dispersion. We treated K562 cells with PEG-CLS. Aggregated cells in solution were dispersed once by pipetting and then were coated with PEG-CLS. We found that cells remained dispersed only when coated with PEG-CLS, which was dependent on the PEG-CLS concentration, otherwise cells reverted to the aggregated form (Figure 3A–C). When the PEG-CLS molecules were washed out, the cells that had been dispersed also rapidly reverted to an aggregated state, as expected (Figure 3B, C). This system enabled the formation of simple and durable cell dispersions, without the addition of digesting proteins to the cells or the removal of aggregated cells. Therefore, this method may be useful for single cell analysis in flow cytometry and genetics.

**Figure 3.**
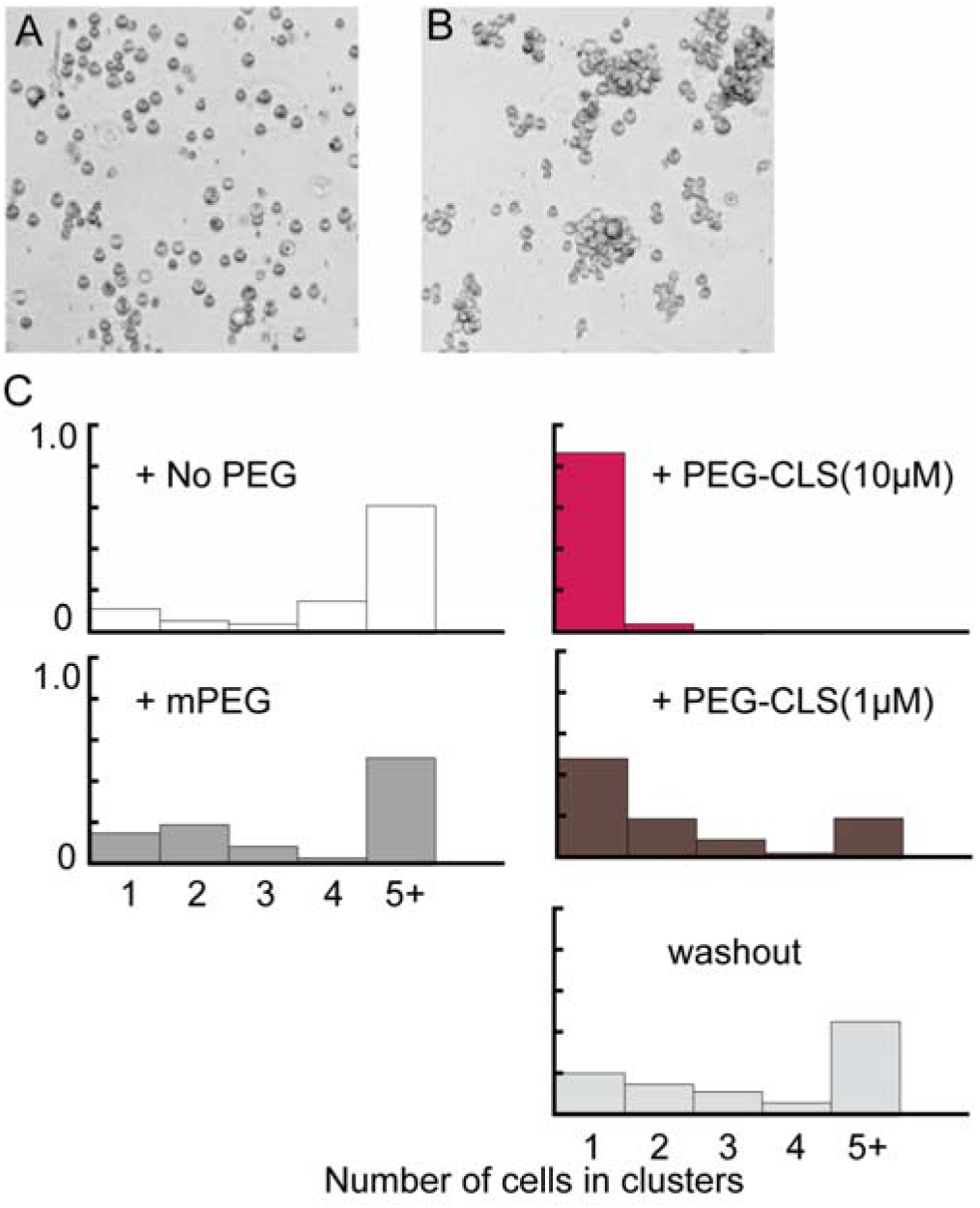
Dispersions of coated cells in solution. (A and B) Images of K562 cells coated with 10 μM 3.4K-PEG-CLS (A) and at 2 h after washout of the PEG-CLS (B). (C) Normalized histogram of the number of K562 cells present as single cells (1) or in clusters of cells (2–4 or 5 or more cells), counted from the images.

We next sought to determine whether coating cells with amphiphilic polymers can regulate cellular processes that take place over a longer time scale, such as cell migration. In a conventional two dimensional migration assay system, cells form anchoring points to a substrate, such as focal adhesions, and convert the internal pushing force generated by polymerized actin filaments against the cell membranes to the motion of whole cell, whereas such anchoring is more complicated in three-dimensional cell migration *in vivo* ^33–34^. The speed of the migrating cells depends on not only the cytoskeleton and membrane structure, but also on the cell–substrate interaction ^35^, thus, many different synthetic and fabricated substrates have been used for the regulation of cell migration ^36–37^.

In contrast to previous studies that used synthetic substrates, we used soluble materials for the modulation of cell migration. We used a cell migration assay, where the number and shape of NIH3T3 cells that migrated toward a prepared gap region on the substrate could be determined. At a concentration of 5 μM of gel-CLS the detachment of NIH3T3 cells from the substrate was not observed. However, we observed an appreciable effect of the cell coating on the migration. There were approximately 50% less cells coated with gel-CLS that migrated to the gap compared with control cells without the coating (Figure 4A–C). Therefore, our results indicated that coating cells with amphiphilic polymers can slow down the cell migration.

**Figure 4.**
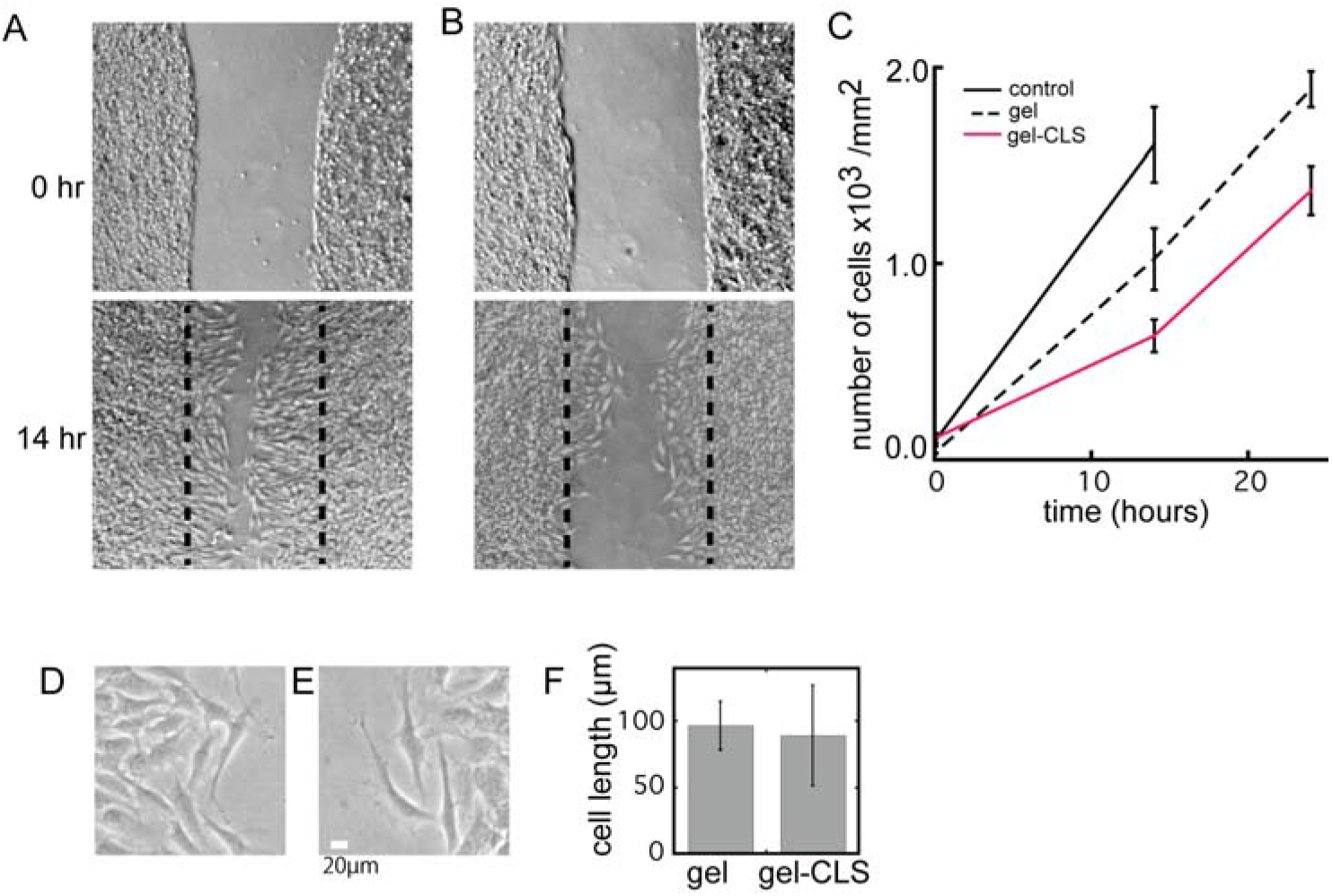
Cell coating can regulate cell migration. (A and B) Images of NIH3T3 cells in wound healing assays at 0 and 14.5 h, where cells were coated with 5 μM of unmodified (A) or cholesterol-modified (B) gelatin. (C) The numbers of cells that migrated toward the gap region on the substrate at different time points, either untreated or treated with unmodified or cholesterol-modified gelatin. (D–F) Magnified view of migrating cells treated with unmodified (D) or cholesterol-modified (E) gelatin, and the average length of the longer axis of these migrating cells (F).

Although there was a difference in the migration speeds, we could not resolve this effect with the changes in cell morphology induced by the cell coating (Figure 4D–F). The results suggested that coating with a low concentration of polymers did not appreciably alter the membrane elastic properties or the internal forces of the cytoskeleton. Instead, we speculated that the membrane coating may partially inhibit and slow down the formation of focal adhesions by inducing steric repulsion between the cells and the substrates, and this may slow down the whole migration process. Further analyses are required to fully understand these processes.

In addition, we investigated whether coating the cells with amphiphilic polymers can also disperse cells in tissues *in vivo*, as was shown *in vitro*. Cell dispersion is critical for collecting and separating cells from tissues, and many strategies, including enzymatic and physical disruption have been developed ^38–40^. Although the existing technologies have been very useful, methods for higher quality and more efficient cell separation from tissues are desirable, particularly for single cell genetic analyses.

As a model system, we tested our strategy in the isolation of infiltrating immune cells in tumors. It is important to obtain high quality samples of these immune cells for analysis, as these cells govern the efficacy of tumor immunology therapies, including checkpoint inhibitor drugs ^6^. We inoculated murine colon adenocarcinoma cells MC-38 in C57BL/6 mice ^41^. After the dissection of isolated tumors, both PEG-CLS and various enzymes were added and the immune cells were separated. The collected cells had good viability (> 90%) and were stained with antibodies against marker proteins for immune cells. We detected increases in the numbers of CD45, CD11b, and CD11c positive cells obtained from PEG-CLS-treated tumors compared with those in the control samples, and the amount of increase was comparable to tumors treated with enzymes (Figure 5A–D). The marker proteins used are expressed in various immune cells: CD45 phosphatase is expressed in all lymphocytes and is weakly expressed in other leukocytes; the integrin CD11b is expressed in natural killer cells, granulocytes, and macrophages; and the integrin CD11c is expressed in monocytes, natural killer cells, and macrophages. Thus, increases in the signals from these proteins indicated an overall increase in the amount of isolated immune cells.

**Figure 5.**
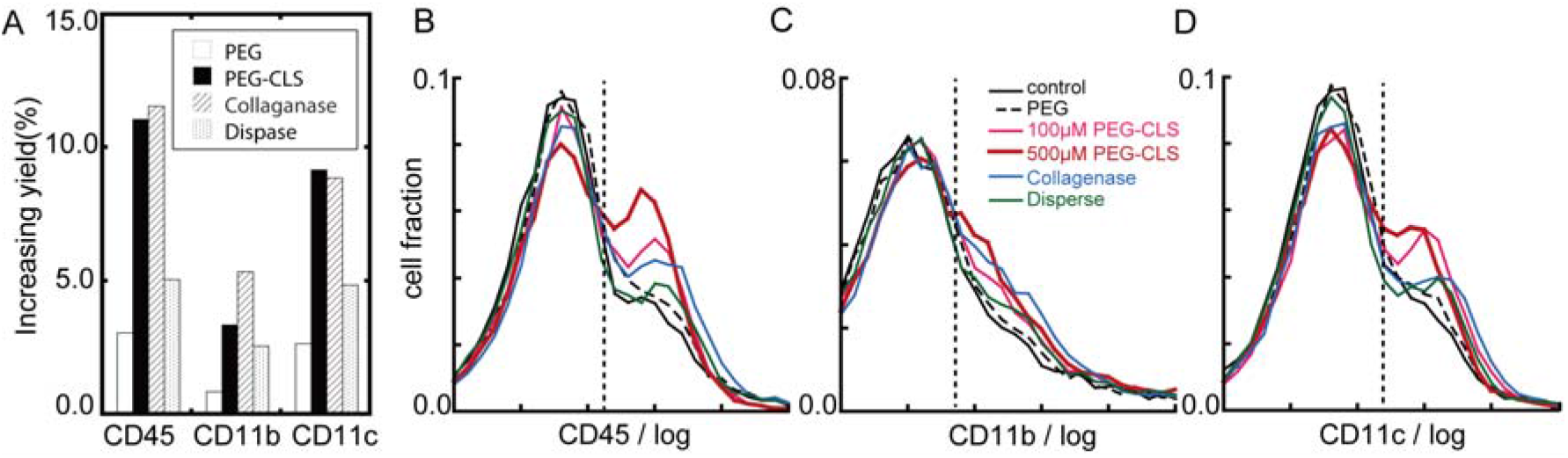
Cell dispersion in tissues. Flow cytometry analysis of tumor infiltrated immune cells extracted from dissected MC38 tumors untreated or treated with mPEG, PEG-CLS at two different concentrations, or the enzymes collagenase-DNAse mix or dispase. Tumor infiltrated immune cells were stained with antibodies against CD45 (B), CD11b (C), and CD11c (D), and the bar graph shows the yields of cells with these marker proteins compared with the untreated control sample (A).

These results suggested that this chemical approach for cell dispersion may be potentially useful for collecting cells from tissues. The coating of cell membranes with amphiphilic polymers is a mild process, as it is reversible as shown in the *in vitro* experiments and does not involve any chemical modification or disruption. In addition, this strategy can be combined with other physical and enzymatic tissue digestion protocols; thus, more sophisticated combination methods could be developed in the future.

## Summary and Conclusions

We have developed a method to chemically control cell adhesion and dispersion by introducing cholesterol-modified polymers as coating materials for cell membranes. We concluded that these coating polymers generated two repulsive forces, tension and steric repulsion. Both tension and steric repulsion are effective for cell detachment, however further analysis is necessary to fully understand how the repulsive forces are generated, which will help optimization of the chemical control of cell adhesion versus dispersion of target cells. We investigated cell detachment, cell dispersion, and cell migration as potential applications for the cell-coating method. However, the repulsive forces, both tension and steric repulsion, are intrinsic in cells as cells express many proteins, and such native repulsive forces may be balanced with other forces to integrate cellular communications. Therefore, further study of this mechanochemical control method may contribute to the future development of native-like, synthetic cell organizational systems.

